# Amyloid pathology impairs experience-dependent inhibitory synaptic plasticity

**DOI:** 10.1101/2023.05.04.539450

**Authors:** Suraj Niraula, Shirley ShiDu Yan, Jaichandar Subramanian

## Abstract

Alzheimer’s disease patients and mouse models exhibit aberrant neuronal activity and altered excitatory-to-inhibitory synaptic ratio. Using multicolor two-photon microscopy, we test how amyloid pathology alters the structural dynamics of excitatory and inhibitory synapses and their adaptation to altered visual experience *in vivo* in the visual cortex. We show that the baseline dynamics of mature excitatory synapses and their adaptation to visual deprivation are not altered in amyloidosis. Likewise, the baseline dynamics of inhibitory synapses are not affected. In contrast, visual deprivation fails to induce inhibitory synapse loss in amyloidosis, a phenomenon observed in nonpathological conditions. Intriguingly, inhibitory synapse loss associated with visual deprivation in nonpathological mice is accompanied by the broadening of spontaneous but not visually evoked calcium transients. However, such broadening does not manifest in the context of amyloidosis. We also show that excitatory and inhibitory synapse loss is locally clustered under the nonpathological state. In contrast, a fraction of synapse loss is not locally clustered in amyloidosis, indicating an impairment in inhibitory synapse adaptation to changes in excitatory synaptic activity.

## Introduction

Coordination of excitation and inhibition governs the activity of cortical neurons and shapes computation (Isaacson and Scanziani, 2011). Neuronal hyperactivity has been observed early in Alzheimer’s disease (AD) patients and mouse models, suggesting impaired neuronal activity homeostasis (Busche and Konnerth, 2016; Palop and Mucke, 2016; Frere and Slutsky, 2018). Interestingly, hyper and hypo-active neurons are present at the same time in mouse models (Busche et al., 2008). We also recently found that a small fraction of neurons in the visual cortex of hAPP mice show higher responsiveness and a larger fraction of neurons are non-responsive to natural image stimuli compared to nonpathological mice (Niraula et al., 2023). Hyper and hypoactive neurons in amyloidosis indicate that changes to activity levels may not trigger compensatory synaptic plasticity mechanisms. This failure may stem from multiple causes, such as impaired detection of excitability changes, or if they are detected, the subsequent response, mediated through synaptic plasticity mechanisms, is impaired (Styr and Slutsky, 2018).

Inhibitory synaptic plasticity is one of the compensatory adaptations to changes in excitatory drive (Gainey and Feldman, 2017; Chen et al., 2022). Postmortem tissues of AD patients exhibit synapse loss, with many studies, but not all, showing a reduction in excitatory or inhibitory synapses (Melgosa-Ecenarro et al., 2023). Contradictory findings regarding the early involvement of inhibitory synapses in AD pathogenesis are also observed in various mouse models of the disease (Melgosa-Ecenarro et al., 2023). It is unclear whether inhibitory synapse plasticity and their adaptation to changes in excitatory drive is altered in AD-associated pathology.

Structural synaptic adaptations to changes in neural activity patterns *in vivo* are typically studied in sensory cortices by altering sensory experience-evoked activity (Gainey and Feldman, 2017; Lee and Kirkwood, 2019). Under nonpathological conditions, sensory deprivation leads to structural plasticity of excitatory and inhibitory synapses, favoring increased excitability of adult cortical neurons *in vivo*. Visual deprivation increases the strength and size of excitatory synapses in the visual cortex and modulates their dynamics (Goel and Lee, 2007; Keck et al., 2008; Hofer et al., 2009; Coleman et al., 2010; Keck et al., 2013; Barnes et al., 2017; Zhou, Lai and Gan, 2017; Sammons, Clopath and Barnes, 2018; Sun et al., 2019). In addition, adult layer 2/3 neurons exhibit increased loss of inhibitory synapses following visual deprivation (Keck et al., 2011; Chen et al., 2012; van Versendaal et al., 2012; Villa et al., 2016; Niraula et al., 2023), and mature excitatory synapses become less dynamic (Subramanian et al., 2019).

*In vivo* multiphoton imaging of structural synaptic dynamics in mouse models of AD thus far is limited to dendritic spines and axonal boutons (Liebscher and Meyer-Luehmann, 2012; Dorostkar et al., 2015; Subramanian, Savage and Tremblay, 2020). These studies reveal increased synaptic dynamics favoring their loss, particularly closer to amyloid plaques, though the phenotype varies depending on the age, mouse strain, or the brain regions examined (Subramanian, Savage and Tremblay, 2020). How amyloid disrupts inhibitory synapses is less understood and remains controversial (Jimenez-Balado and Eich, 2021; Melgosa-Ecenarro et al., 2023). Though amyloid accumulates in the boutons of inhibitory neurons, the basal structural dynamics of these boutons are not significantly altered (Ruiter et al., 2021). However, some studies but not others have found axonal degeneration of inhibitory neurons and structural synapse loss (Garcia-Marin et al., 2009; Palop, Mucke and Roberson, 2011; Mitew et al., 2013; Kiss et al., 2016; Schmid et al., 2016; Umeda et al., 2017; Hollnagel et al., 2019; Petrache et al., 2019; Ruiter, Herstel and Wierenga, 2020; Shimojo et al., 2020; Sos et al., 2020; Montero-Crespo et al., 2021; Kurucu et al., 2022; Niraula et al., 2023; Scaduto et al., 2023). A challenge to studying inhibitory postsynaptic dynamics *in vivo* is that they are primarily present in the dendritic shaft of excitatory neurons and lack a morphological surrogate, such as dendritic spines, which are typically used as a proxy for excitatory synapses. However, most dynamic spines carry immature excitatory synapses (Villa et al., 2016; Vardalaki, Chung and Harnett, 2022); therefore, how amyloid influences mature excitatory synapse dynamics *in vivo* remains unclear. Whether amyloid pathology disrupts the experience-dependent structural plasticity of mature excitatory or inhibitory synapses *in vivo* is unclear.

Using a synaptic labeling approach that reliably detects mature excitatory and inhibitory synapses of cortical neurons and *in vivo* multicolor two-photon imaging, we show that the baseline dynamics of mature excitatory and inhibitory synapses are not altered in the visual cortex of a mouse model of AD (hAPP mice - J20 line (Harris et al., 2010)). Interestingly, visual deprivation-evoked structural loss of inhibitory synapses is disrupted in these mice. In contrast, reduced excitatory synaptic dynamics associated with visual deprivation are preserved, indicating that neurons in amyloid pathology retain their ability to sense changes in experience-evoked excitability. Accompanying inhibitory synapse loss, we also observe the broadening of spontaneous but not visually evoked calcium transients selectively in nonpathological mice. We show that impaired structural adaptation of inhibitory synapses following visual deprivation could stem from the impaired local coupling of excitation and inhibition, manifested as a reduced spatial clustering of the loss of excitatory and inhibitory synapses.

## Methods

### Mice

The University of Kansas Institute of Animal Use and Care Committee has authorized all animal procedures and complies with the NIH standards for the use and care of vertebrate animals. Transgenic hAPP mice overexpressing a mutant human form of amyloid precursor protein (APP) that encodes hAPP695, hAPP751, and hAPP770 bearing mutations linked to familial AD [APPV717F (Indiana), KM670/671NL(Swedish)], J-20 line were obtained from Jackson Laboratory). J20 mice and wild-type (WT) females from the same background were bred to generate heterozygotes for the hAPP transgene. J20 males and C57BL/6J-Tg (Thy1-GCaMP6s) GP4.3Dkim/J (Strain: 024275, JAX; (Chen et al., 2013)) females were bred to generate J20-GCaMP6s mice. Five mice at most were kept in a cage, but following cranial window surgery, they were individually housed on a 12h light/12h dark cycle or 24 hours darkness for visual deprivation experiments. Both genders were used in the study.

### DNA constructs

The Cre-dependent TdTomato (pFudioTdTomatoW), Teal-gephyrin (pFudioTealgephyrinW), and PSD-95-venus (pFudioPSD-95venusW) plasmids were a kind gift from Dr. Elly Nedivi. The pSIN-W-PGK-Cre plasmid is used to express Cre recombinase (Subramanian, Dye and Morozov, 2013). The Cre-dependent expression of fluorescently labeled synaptic markers PSD-95 and gephyrin have been shown to accurately represent excitatory and inhibitory synapses, respectively (Villa et al., 2016).

### In-utero electroporation (IUE)

Timed pregnancies were established between heterozygous J20 males and WT females with the same genetic background. Half of the litter was heterozygous for the APP transgene, while the other half was WT (control). Using a 32-gauge Hamilton Syringe (Hamilton company), plasmids diluted in 1 µl of Tris-EDTA (1:1:0.5:0.15 molar ratios of pFudioTdTomatoW, pFudioTealgephyrinW, pFudioPSD-95venusW, and pSIN-W-PGK-Cre, respectively) were injected in the lateral ventricle of E15.5 to E16.5 embryos. A square wave electroporator (ECM830, Harvard Apparatus) was used to deliver five pulses of 36 V (50 ms duration at 1 Hz) to a pair of platinum electrodes (Protech International) targeted at the visual cortex.

### Cranial window

A cranial window was placed in 4-6-month-old J20 and WT mice over the visual cortex in the right hemisphere. An incision was made above the midline of the skull. The pericranium was softly scraped, and soft tissues were reflected laterally by blunt dissection. A biopsy punch was used to score a 5-mm-diameter circle covering the visual cortex. Using a fine drill and a sterile 0.5mm diameter round burr (Fine Science Tools), the skull was thinned along the scored circle. Fine forceps were used to carefully remove the bone flap, leaving the dura intact. A sterile, 5-mm diameter circular glass coverslip (Harvard Apparatus) was placed over the opening. To secure the coverslips in place, firm pressure was applied while Vetbond was placed over the area where the coverslip and bone met. Over the exposed skull, Metabond (C&B Metabond) was placed and a titanium head post was attached to the window about two weeks following the surgery.

### Viral injections

For GCaMP6 expression, we used pAAV.syn.GCamMP6s.WPRE.SV40 (viral titer/ml: 1.3 x 10^13, Addgene 100843-AAV9). For jRGECO and GPHN.FingR.EGFP expression, we used pAAV.Syn.NES-jRGECO1a.WPRE.SV40 (viral titer/ml: 3.2 x 10^13, Addgene 100854-AAV9), pCAG.DIO.GPHN.FingR.EGFP2F (viral titer/ml: 2.4 x 10^13; a kind gift from Don Arnold) and pENN.AAV.CamKII 0.4.Cre.SV40 (viral titer/ml: 2.3 x 10^13 GC/ml (Addgene 105558-AAV9) in the ratio 2:1:1, respectively. We injected the viruses in the right primary visual cortex in three locations centered on stereotactic coordinates 2.9 mm lateral and 0.5 mm anterior to lambda. We injected at a depth of 150-300 µm from the dura and 150 nl per injection site at the rate of 25-30 nl/minute. The needle was slowly retracted after 5 minutes of the injection on each site, and a cranial window encompassing the viral injection sites was placed.

### Optical intrinsic signal imaging

∼14 days after cranial window surgeries, optical intrinsic signal imaging was performed to map the location of the visual cortex. A custom-designed upright microscope with a 4X objective (Nikon) was used for imaging. Mice were lightly sedated using isoflurane and positioned 20 cm in front of a high refresh rate monitor showing a horizontal bar drifting at 10 Hz. Images were captured at 5Hz with an sCMOS camera (1024 x 1024 pixels; Photometrics). The cortex was illuminated (500–600 m below the dura) using 610 nm light delivered by a fiber-coupled LED controlled by T-Cube LED drivers (Thorlabs). A 470 nm light was used to image reference vasculature. Cortical intrinsic signals were computed by extracting the Fourier component of light reflectance changes to matched stimulus frequency from downsized images (256×256 pixels). The magnitude maps were thresholded at 30% of the peak response amplitude. The fractional change in reflectance represents response magnitude. The magnitude maps were superimposed over the 470nm reference image to map the visual cortex.

### Widefield calcium imaging

Instead of using intrinsic signal imaging to map the location of the visual cortex in GCaMP6s transgenic mice, widefield calcium imaging was employed. The mapping methodology was comparable to intrinsic signal imaging, except fluorescence was imaged as opposed to reflected light. GCaMP6 was excited by an LED (Lambda FLED, Sutter) filtered through a bandpass filter (470/40, 49002 Chroma), and the emission was filtered with a 525/50 bandpass filter.

### Two-photon imaging

Synaptic structural imaging was performed on isoflurane anesthetized mice with sparsely labeled neurons in the mapped visual cortex using a Sutter MOM multiphoton microscope. The Ti: Sapphire laser (MaiTai HP: Newport SpectraPhysics; 915 nm) was directed toward the microscope using table optics. A polarizing beam splitter and a rotating half-wave plate were used to regulate laser power. A pair of galvanometric mirrors scan the laser beams to the back aperture of the objective (Nikon 16X 0.8 NA). The output power from the objective was set to 40-50 mW. The same objective was used to gather the emission signal, which was then routed through appropriate bandpass filters (488/50, 540/50, and 617/73 for Teal, YFP, and TdTomato fluorescence, respectively) and three GaASP PMTs. Image acquisition was controlled by ScanImage (Vidrio Technologies), and images were obtained at 0.16Hz. The imaging field covered 133×133x∼150 μm (1024 x 1024 XY pixels, Z step - 1 μm).

For measuring calcium transients during visual deprivation, transgenic (3 WT, 2 hAPP mice) and virally delivered GCaMP6s (2 WT, 2 hAPP mice) were used. Neurons within the mapped visual cortex (∼100-150 μm below the dura) were imaged at 4.22 Hz in head-restrained awake mice restrained in a body tube. The excitation wavelength was set to 940 nm, and the power was adjusted to avoid signal saturation. The imaging field was a single Z frame of 336 x 336 um (256 x 256 pixels) consisting of ∼50-100 cells. In each imaging session, we first imaged calcium transients in complete darkness (for spontaneous activity; 213 seconds), followed by visual stimuli (415 seconds). Visual stimuli consisted of six trials of three seconds of phase reversing orientation grating stimuli (0°, 30°, 60°, 90°, 120°, 150°) that do not drift and natural images. Six seconds of gray screen interspersed grating and natural image stimuli were presented in random order in each trial.

jRGECO imaging was restricted to spontaneous activity imaging and was performed similarly to GCaMP6 imaging, except that the excitation wavelength was set to 1040 nm. Immediately following jRGECO imaging, mice were anesthetized and imaged for GPHN.FingR-GFP. The imaging field for gephyrin FingR-GFP covered 133×133×25 μm (1024 x 1024 XY pixels, Z step – 0.25 μm). The excitation wavelength was set to 915 nm. The power was adjusted to avoid signal saturation.

### *In vivo* synaptic imaging analysis

The signal collected in each PMT (channel) is a combination of signals from the three fluorophores (Teal, Venus, and TdTomato) due to their overlapping emission spectra. We used spectral linear unmixing to reassign the signal from each fluorophore to the appropriate channel. Each image consisted of three channels: cell fill (TdTomato), PSD-95 (Venus), and Gephyrin (Teal) channels. First, gephyrin and PSD-95 puncta were marked if they were present in two consecutive frames and consisted of at least 8-9 or 4-5 clustered pixels, respectively, in mean-filtered, volume-corrected images. For volume correction (normalization of the signal relative to local dendritic volume), we normalized the fluorescence in the synaptic channels to that of the cell fill channel. A normalization factor was calculated as the ratio of the mean pixel value of a chosen dendrite in the cell fill channel to the synaptic channel. Each pixel value in the synaptic channel was then multiplied by the normalization factor, and the pixel value of the cell fill channel was subtracted on a pixel-to-pixel basis. A custom-written 4D point tracking system implemented in Fiji using a modified version of the ObjectJ plugin (Villa et al., 2016) was used to transfer labeled markers (indicating each synapse type or no synapse) to matched locations on images from subsequent imaging sessions. Synaptic markers were transferred back to the identical location on the unmixed image for quantification. A custom-written macro was used to place a 5×5 pixel box (synaptic region of interest (ROI)) at the center of synaptic puncta for PSD-95 on the spines and shaft gephyrin. The box overlapped with part of both puncta for dually innervated spines containing both PSD-95 and gephyrin puncta. 5×5 pixel boxes (background ROI) were also placed on dendritic shaft locations lacking visible puncta to calculate background fluorescence. The background ROI boxes, equaling the number of identified synapses, were placed along the entire length of the dendritic segment used for synapse identification. PSD-95 puncta on spines were classified as excitatory synapses based on a clustering index (CI_PSD-95_), which is calculated as

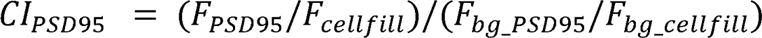

where F_PSD-95_ and F_cellfill_ are the mean fluorescence of synaptic ROI from PSD-95 and cell fill channel of an identified puncta, respectively, and F_bg_PSD-95_ and _Fbg_cellfill_ are mean + 3x standard deviation of fluorescence of ten nearest background ROIs to that puncta. Similarly, CI_gephyrin_ is calculated as

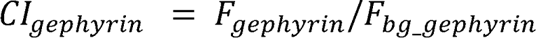

where F_gephyrin_ and F_cellfill_ are the mean fluorescence of synaptic ROI and background ROI of the gephyrin channel, respectively. PSD-95 and gephyrin are considered excitatory and inhibitory synapses if their clustering index is greater than one.

The gain and loss of a specific type of synapse between sessions were determined by calculating the number of newly formed puncta or lost puncta of that synapse type and dividing it by the total number of puncta of that synapse type in the later or previous session, respectively.

For nearest neighbor distance analysis, we first calculated the cumulative distance along the trace of the dendrite generated by Simple Neurite Tracer (SNT) by iteratively summing the Euclidean distances between consecutive points (XYZ coordinates) of the SNT trace. This creates a continuous distance measurement along the SNT trace. We then computed pairwise distances between each synapse marker and all points along the SNT trace using the Euclidean distance formula, considering the 3D coordinates (X, Y, Z) of each marker and trace point. For each synapse marker, we identified the SNT trace point that is closest to it by finding the minimum distance from the pairwise distance calculations using the ‘pdist2’ function. The distance of a synaptic puncta from the soma was calculated by summation of the distance of the puncta to the center of the nearest dendritic shaft and the distance from that point to the soma. To identify the nearest synapse, the distances of all synapses to each other were first calculated, and the nearest synapse was determined using the ‘knnsearch’ function in MATLAB.

The nearest neighbor of both excitatory and inhibitory dynamic synapses was calculated using a similar approach. To evaluate whether the observed spatial relationships between synapse losses on dendritic branches deviated from a random distribution, we used experimentally observed data on synaptic density, dendritic length, and the count of lost synapses for each dendritic segment. We also used the actual dendritic locations of “source” synapse losses, which act as reference points for identifying the closest “target” synapse losses. We then generated random distributions of target synapse losses and calculated the nearest neighbor distance. This process was repeated for each source synapse across all dendrites and averaged. We iterated it 10,000 times to generate a random distribution of nearest neighbor distances of lost target synapses to lost source synapses.

We tracked 2453 PSD-95^+^ spines in WT mice, 2156 in hAPP mice, 1111 gephyrin puncta in WT, and 1073 in hAPP mice across three imaging sessions. These structures were identified in 54 dendrites spanning a length of 4133 µm in WT mice and 48 dendrites spanning 3719 µm in hAPP mice. The synaptic density from the first imaging session for some of the analyzed hAPP neurons has been published (Niraula et al., 2023).

The objectJ plugin described above was also used to track gephyrin puncta identified by GPHN.FingR-GFP across sessions. A 5×5 ROI was placed over the identified puncta, and two additional ROIs were placed in the nearby background using a custom-written macro. Puncta were scored as gephyrin if the fluorescence was greater than three standard deviations from the mean of all background ROIs on the image and if the puncta was present in 5 consecutive imaging frames (0.25 µm/frame). We tracked 422 puncta from 6 WT mice.

### Calcium imaging analysis

The data (Figure 4-C) on spontaneous calcium imaging is an independent analysis from the same raw data used for other analyses previously published (Niraula et al., 2023). Suite2p was used to perform motion registration and ROI detection on the time-series images (Pachitariu et al., 2017) with tau set at 2 seconds (GCaMP6s) or 0.7 (for jRGECO). If the soma was discernible in the mean or maximum projection picture, ROIs generated by Suite2p were selected as cells (cellular ROI). The neuropil-corrected fluorescence (Fcorr) is calculated as F - (0.5xFneu). dF/F0 is calculated as (Fcorr – F0)/F0, where F0 is defined as the mode of the Fcorr density distribution. For each neuron, the mean dF/F0 over every three-second period was calculated for 228 seconds (76 x 3 seconds) and averaged. The percentage of neurons in each indicated dF/F0 bin was calculated as the number of neurons in that bin divided by the total number of identified neurons for each mouse. To assess functional connectivity in cellular ROIs obtained from Suite2p, we deconvolved spikes and applied a threshold (>2 SD from the mean) before binarizing the data. We then determined functional connections between pairs of neurons by comparing their coactive frames to a distribution generated from 1000 random circular shifts of their activity. Neuron pairs with coactive frames exceeding the 95th percentile of this distribution were considered functionally connected. We used the resulting functional connectivity matrix to determine the node degree of each neuron, which represents the number of edges (or connections) connected to that node over the entire imaging period. We calculated node degrees using MATLAB’s graph and degree functions.

To assess the area under each transient’s curve (AUC), we set a threshold of mean plus two standard deviations of the entire spontaneous or visually evoked stimulus periods. We used MATLAB’s ‘findpeaks’ function to identify peaks, and the area under each peak above the threshold was calculated using the ‘trapz’ function. The area of all the transients during the spontaneous imaging period was averaged for each neuron. We excluded the gray screen imaging frames from analysis for visually evoked calcium transients and averaged all transients across different stimuli for each neuron. Neurons were not matched between imaging sessions.

### Statistics

Statistics were performed using MATLAB, SPSS, or GraphPad Prism. P-values and statistical procedures are provided in the figure legends. P<0.05 is considered significant unless mentioned otherwise (for Bonferroni corrections). To compare the mean of the observed nearest distances of target and source synapse loss with that of randomly generated distribution, we obtained a left-sided P value as the mean was always to the left of the random distribution.

## Results

### Selective disruption of homeostatic structural plasticity of inhibitory synapses on the dendritic shaft in amyloid pathology

To study how amyloid pathology influences the dynamics of excitatory and inhibitory synapses *in vivo*, we used an in-utero electroporation based synaptic labeling strategy that sparsely labels layer 2/3 cortical neurons in the visual cortex (Chen et al., 2012; Villa et al., 2016; Subramanian et al., 2019). Three fluorescent proteins, TdTomato (cell fill), PSD-95-venus (excitatory synaptic marker), and Teal-Gephyrin (inhibitory synaptic marker), were expressed in a Cre recombinase dependent manner (Figure 1A). Previous studies have shown that this approach reliably represents excitatory and inhibitory synapses in the same neurons *in vivo* along with dendritic spines (Chen et al., 2012; Villa et al., 2016). We used multicolor two-photon microscopy to image individual neurons expressing all fluorescent markers in the visual cortex *in vivo* under normal and visual deprivation conditions. Visual deprivation allows us to examine structural synaptic adaptation to reduced experience-evoked activity. To assess synaptic dynamics under baseline and altered experience conditions, we imaged the same neurons and synapses thrice with an interval of seven days between imaging sessions (Figure 1A, B). Between the first two imaging sessions, the mice were housed in a 12-hour light/12-hour dark cycle (baseline). Immediately following the second imaging session, mice were transferred to 24-hour darkness until the final imaging session (visual deprivation). The appearance of new synapses (gain) and disappearance of pre-existing synapses (loss) between the first two imaging sessions represents baseline dynamics, and between the second and third sessions includes changes associated with altered experience (Figure 1C-G).

**Figure 1.**
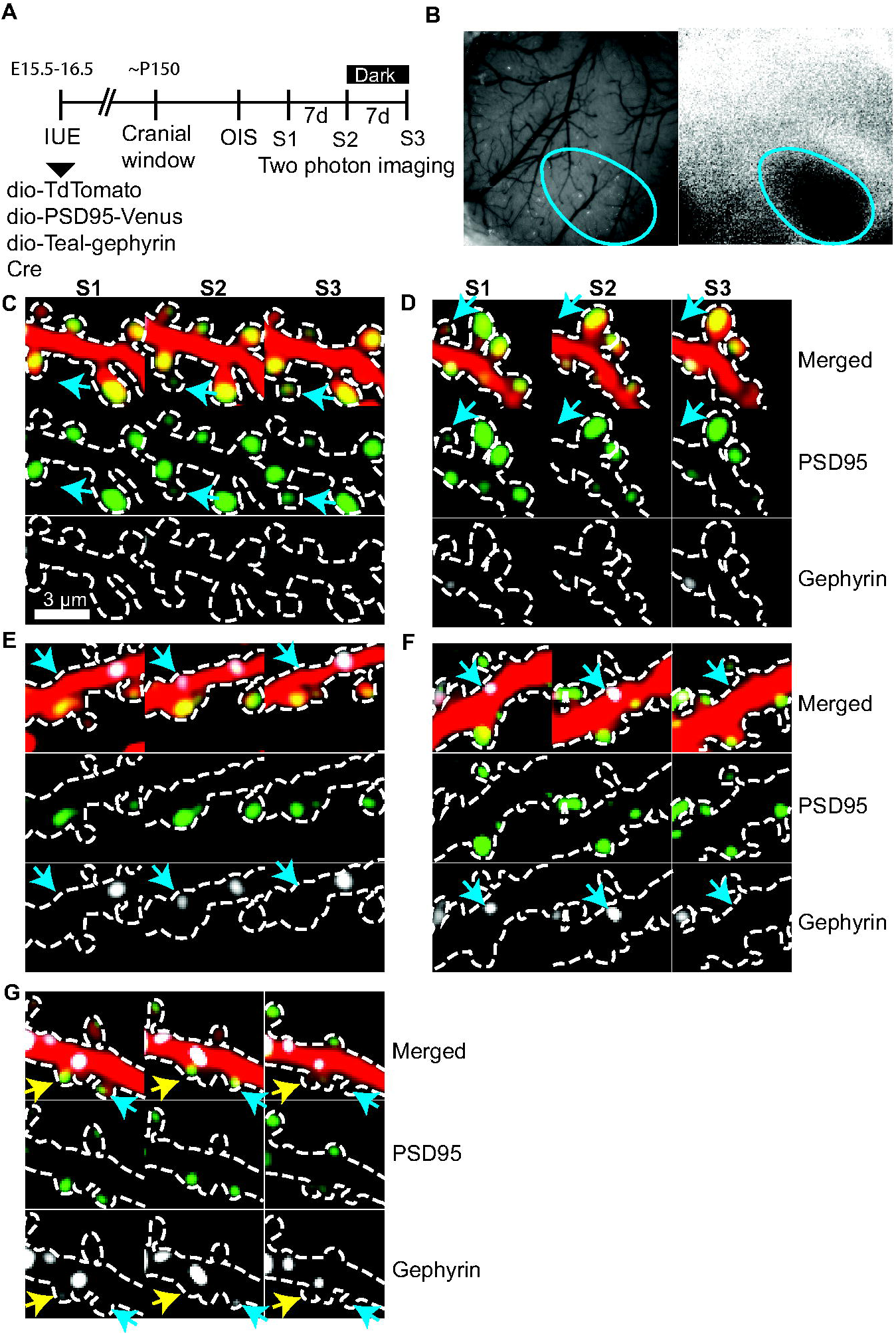
*In vivo* structural imaging of homeostatic synaptic plasticity. **A.** Timeline of imaging. In utero electroporation (IUE) of plasmids expressing Cre recombinase and Cre-dependent markers was performed on embryonic days 15.5-16.5. A cranial window was placed at ∼ five months old, and two weeks later, optical intrinsic signal (OIS) imaging was performed to map the visual cortex. Two-photon imaging of neurons in the mapped visual cortex was performed in three imaging sessions (S1, S2, and S3). Each imaging session was separated by one week, and mice were housed in 24-hour darkness (24 D) between S2 and S3. At all other times, they were housed under 12-hour light: 12-hour dark (12L:12D) cycle. **B.** A 5mm cranial window (left) and a magnitude map (blue oval) of intrinsic signal (right). **C-G.** Pseudocolored images of the same dendritic segments from the three sessions (red: cell fill; green: PSD-95; gray: Gephyrin). PSD-95 and gephyrin channels are shown below the merged images. Gain and loss (blue arrows (C-F) of excitatory synapses (gain – C and loss – D), inhibitory synapse on shaft (gain – E and loss – F), and inhibitory synapses on spines (blue arrow - gain in S1 loss in S2, yellow arrow – loss, G).

The synaptic labeling scheme allows us to detect four types of synaptic structures – dendritic spines with (PSD-95^+^ spines) or without (PSD-95^-^ spines) PSD-95, gephyrin on dendritic shaft (inhibitory shaft synapse), and gephyrin on PSD-95^+^ spines (inhibitory spine synapse). To assess the dynamics of excitatory synapses and inhibitory synapses, we tracked the gain and loss of PSD-95^+^ spines (2453 (WT), 2156 (hAPP)) and gephyrin on the shaft and spines (1111 (WT), 1073 (hAPP)). The average excitatory (wild type (WT): 0.59 ± 0.04/µm; hAPP: 0.58 ± 0.04/µm) and inhibitory (WT: 0.27 ± 0.06/µm; hAPP: 0.28 ± 0.05/µm) synapse densities did not differ between the genotypes.

Under baseline conditions, the gain and loss of all synapse types were balanced. ∼2-3% of excitatory synapses were gained, and a similar fraction was lost during baseline conditions (Figure 2A) for both WT and hAPP mice, indicating that amyloid pathology in the visual cortex prior to plaque does not lead to increased loss of excitatory synapses. We found that visual deprivation reduced the gain of excitatory synapses in WT mice. A similar reduction in the gain of these synapses was also seen in hAPP mice (Figure 2A). Overall, we observed a significant effect for light deprivation (p<0.01; F (1, 13) = 15.06; two-way mixed model ANOVA), but the effect was not significantly different between genotypes or the interaction of light and genotype. In contrast to the gain of excitatory synapses, the loss between two sessions did not differ significantly between genotypes or light conditions (remained between 2-3% across conditions; Figure 2A). These results show that excitatory neurons adapt similarly to changes in visually-evoked excitability under amyloid and nonpathological conditions.

**Figure 2.**
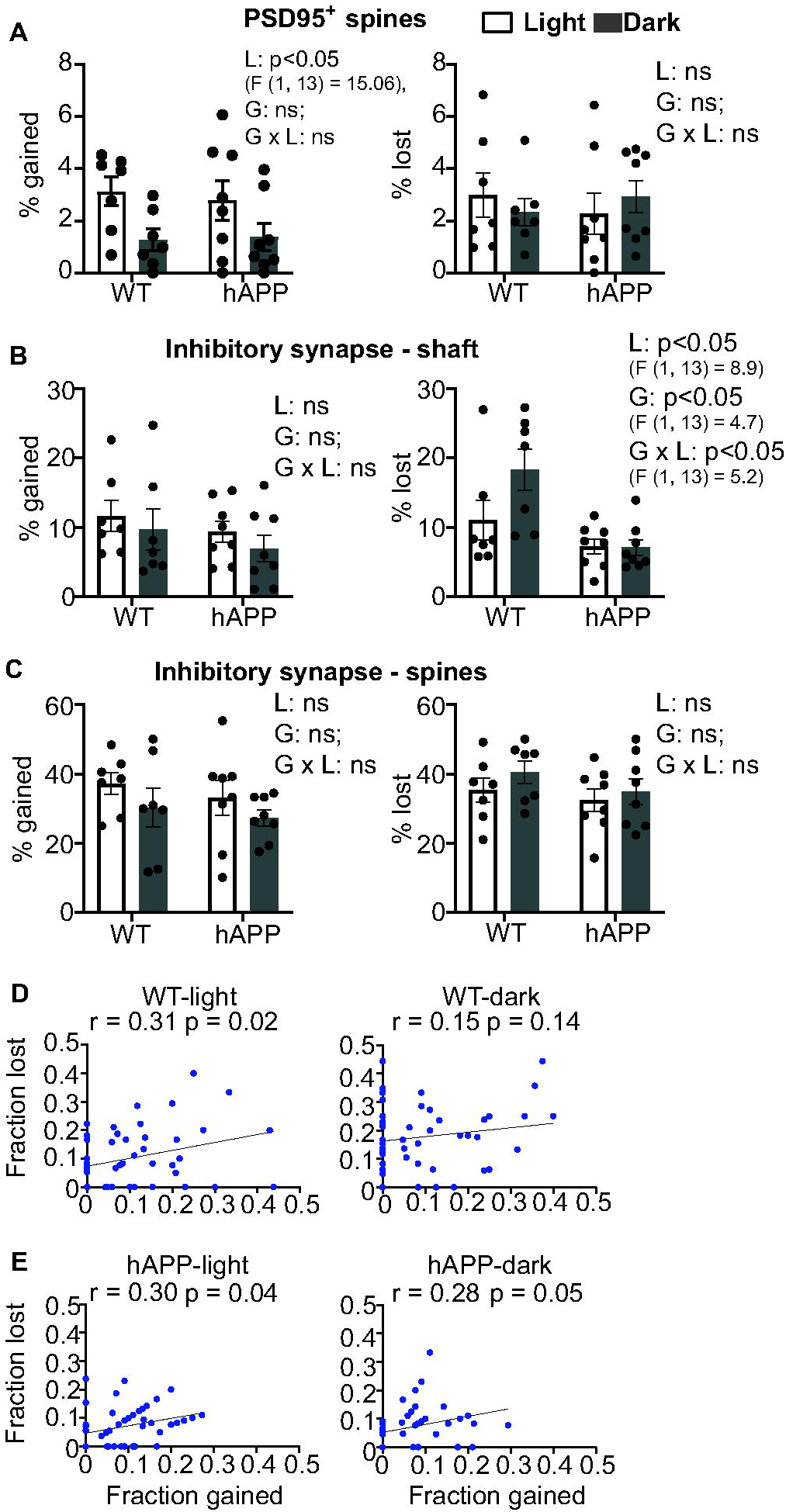
Inhibitory but not excitatory synaptic structural adaptation deficits in amyloid pathology. **A-C.** Percentage of excitatory synapses (A), inhibitory synapses on shaft (B), and inhibitory synapses on spines (C) gained (left) or lost (right) in baseline (light) and visual deprivation (dark) conditions in wild type (WT) and hAPP mice. Circles represent individual neuron values. n = 7 neurons (5 WT mice), 8 neurons (7 hAPP mice). Two-way mixed model ANOVA was used to test the effect of light vs. dark (L), genotype (G), and their interaction (G X L). ns: not significant. Data are presented as mean ± SEM. **D-E**. Correlation of percentage gain and loss of inhibitory shaft synapses on individual dendrites in WT (D) and hAPP (E) mice in baseline and visual deprivation conditions n = 54 dendrites (WT) and 48 dendrites (hAPP).

Inhibitory synapses are more dynamic than excitatory synapses (Villa et al., 2016). Consistently, we found that ∼11% of inhibitory shaft synapses were gained or lost under baseline conditions in WT mice (Figure 2B). The gain and loss were slightly lower in hAPP mice under baseline conditions (∼9% gain and ∼7% loss; Figure 2B), although it did not reach statistical significance. When visually-evoked activity is reduced by visual deprivation, the gain of new inhibitory shaft synapses was reduced by 2-3% in both WT and hAPP mice. However, the effect is not significantly different between the genotypes, light conditions, or their interaction, indicating that pre-plaque amyloid pathology under baseline conditions does not affect the gain of new inhibitory shaft synapses.

Visual deprivation elicits disinhibition as a homeostatic adaptation under nonpathological conditions (Gainey and Feldman, 2017). At the structural level, the loss of inhibitory synapses is increased by visual deprivation in nonpathological mice (Chen et al., 2012; van Versendaal et al., 2012). Consistently, we found that dark adaptation increased the loss of inhibitory shaft synapses by 63% in WT mice (18% loss; Figure 2B). Interestingly, the loss of inhibitory shaft synapses remained identical (∼7% loss) to baseline conditions in hAPP mice (Figure 2B). Consequently, for inhibitory shaft synapses, we found a significant effect for genotype (p<0.05, F (1, 13) = 8.918), visual deprivation (p<0.05, F (1, 13) = 4.733), and their interaction (p<0.05, F (1, 13) = 5.215; two-way mixed model ANOVA; Figure 2B). Inhibitory synapses on spines are more dynamic than inhibitory shaft synapses (Chen et al., 2012; van Versendaal et al., 2012). Consistent with a previous study (Villa et al., 2016), we found that 30-40% of inhibitory synapses on spines were gained or lost under baseline conditions in both WT and hAPP mice; Figure 2C. Though visual deprivation increased the net loss of inhibitory spine synapses (gain/loss: 1.05 (light), 0.75 (dark)) in WT mice, the effect was not different between genotypes, visual deprivation, or their interaction (Figure 2C). These results show that amyloid pathology selectively disrupts the structural adaptation of inhibitory shaft synapses to loss of visual experience.

Since the average gain and loss of inhibitory shaft synapses were balanced under baseline conditions (gain/loss: 1.05 (light)), we tested whether this balance persisted at the level of individual dendrites. We found that the correlation between the gain and loss of inhibitory synapses on dendritic branches was significant in WT mice under baseline conditions (Figure 2D). The correlation reduced during visual deprivation and was no longer significant (Figure 2D). However, the difference in correlation between baseline and visual deprivation conditions was not significantly different (p>0.05, Fisher r to z transformation). These results indicate that the gain and loss of inhibitory shaft synapses are balanced, and visual deprivation mildly disrupts this balance. The correlation between gain and loss remained close to significance in hAPP mice under both baseline and visual deprivation conditions, suggesting that amyloid does not disrupt the balance of gain and loss of inhibitory shaft synapses (Figure 2E).

Different inhibitory neuron subtypes project to various dendritic locations. Parvalbumin-expressing interneurons innervate perisomatic and proximal (<40µm from the soma) dendrites, whereas somatostatin-expressing neurons primarily project to distal dendrites (>40 µm from the soma) (Di Cristo et al., 2004). Therefore, we examined whether visual deprivation associated increases in the loss of inhibitory shaft synapses occur differentially depending on dendritic location, with some locations exhibiting higher synapse loss (hotspots). The density of inhibitory shaft synapses is the highest close to the soma and progressively decreases both in WT and hAPP mice (Figure 3A, B). Visual deprivation uniformly and subtly reduces the density of inhibitory shaft synapses at different distances from the soma in WT mice (Figure 3A). In hAPP mice, the density remained identical across all three sessions for all distances from the soma (Figure 3A). Similarly, the increased loss of inhibitory shaft synapses in visual deprivation is evident at all distances from the soma in WT mice (Figure 3A), whereas the loss of synapses was very similar in hAPP mice at all measured distances from the soma (Figure 3A). These results indicate that there may be no dendritic hotspot for elevated inhibitory synapse loss.

**Figure 3.**
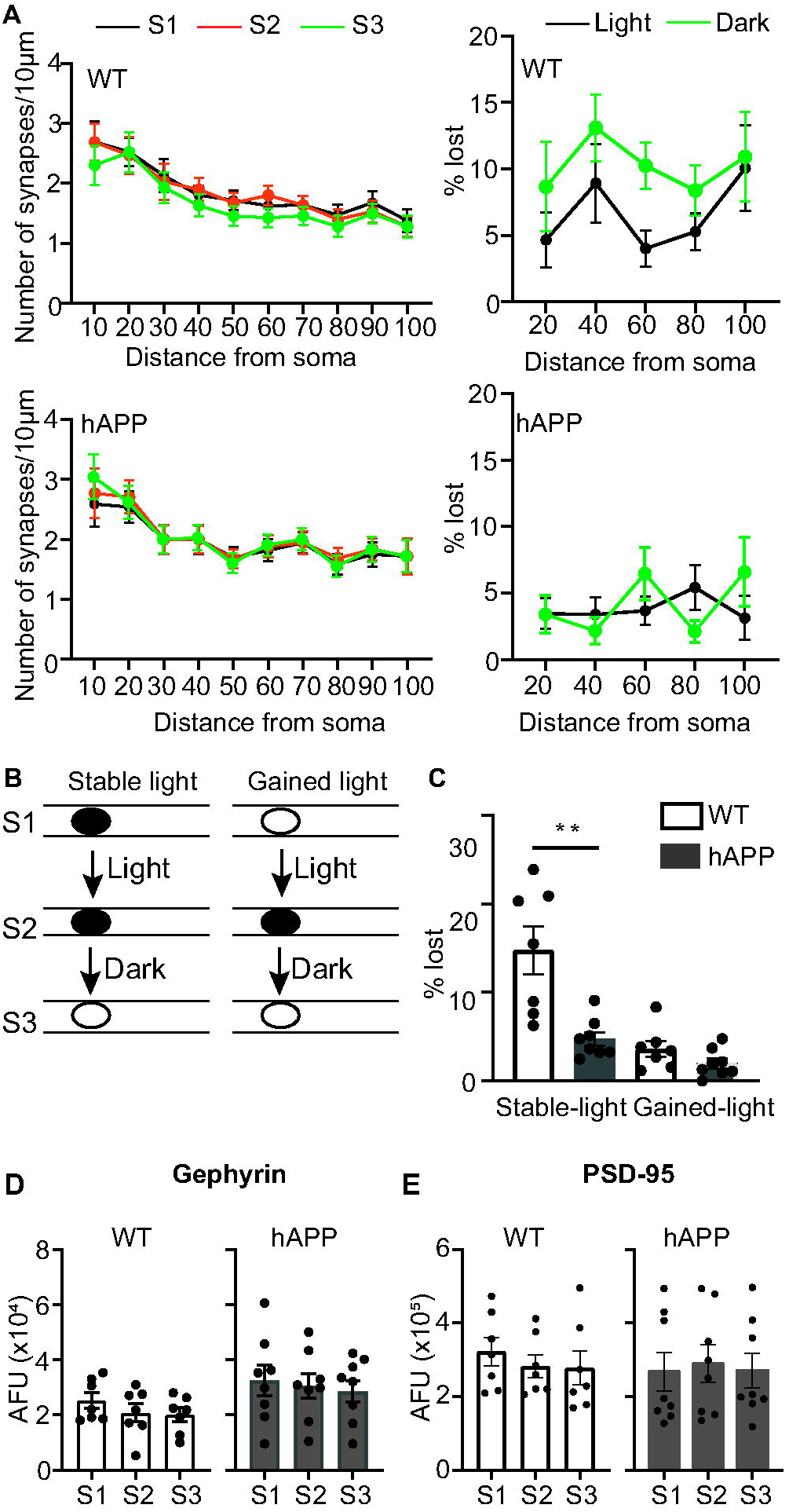
Characterization of dendritic shaft structural disinhibition. **A.** (Left) Density of inhibitory shaft synapses at different distances from soma from the three imaging sessions (S1, S2, and S3) in wild-type (WT) and hAPP mice. (Right) Percentage loss of inhibitory shaft synapses at different distances from the soma in baseline (light) and visual deprivation (dark) conditions in WT and hAPP mice. n = 54 dendrites (WT) and 48 dendrites (hAPP). **B**. Representation of an inhibitory shaft synapse that was present in the first two imaging sessions (stable-light) but was lost in the third session (left) and a synapse that appeared in the second imaging session (gained-light) and disappeared (right). **C**. percentage of stable and gained inhibitory shaft synapses lost following visual deprivation (dark). n = 7 (WT), and 8 (hAPP) neurons. ** p<0.01, *F*_(2,12)_ = 7.2; one-way MANOVA. **D-E.** Average fluorescence (arbitrary fluorescence units (AFU)) of the inhibitory shaft (D) and mature excitatory (PSD-95; E) synapses in the three sessions in WT and hAPP mice. Circles represent neuron values, and data are presented as mean ± SEM.

A fraction of inhibitory synapses tend to appear and disappear at the same dendritic locations (Villa et al., 2016). To test whether the structural loss of inhibitory shaft synapses during visual deprivation was driven mainly by the loss of newly acquired or those present in both imaging sessions, we compared their relative contribution to the total inhibitory shaft synapse loss (Figure 3B). Of the ∼18% synapses that were lost in WT mice in the dark, only 3.6% of them were newly formed synapses in the second session, and the rest (14.8%) were synapses present in the first and second sessions (Figure 3C). In hAPP mice, of the 7% of shaft synapses that were lost during visual deprivation, ∼5% were present in both sessions, and 2% were newly gained synapses (Figure 3C). The loss of newly gained synapses did not differ significantly; therefore, the synapses present in the first two sessions were more stable in hAPP mice (Figure 3C).

Since the size of gephyrin puncta correlates with the synaptic strength (Villa et al., 2016), we tested whether visual deprivation decreases the average fluorescence intensity of shaft synapses that remained stable across the three sessions. We found no difference in average gephyrin fluorescence between sessions in WT and hAPP mice (Figure 3D). Visual deprivation increases dendritic spine size or gain in layer 5 neurons (Hofer et al., 2009; Keck et al., 2013; Barnes et al., 2015), indicating increased excitatory synaptic strength. To test whether excitatory synapse strength is altered by visual deprivation, we compared whether stable dendritic spines show changes in PSD-95 fluorescence, which correlates with synaptic strength (Fortin et al., 2014), following dark exposure. Consistent with studies that found no difference in spine size or gain in layer 2/3 neurons upon visual deprivation (Hofer et al., 2009; Barnes et al., 2015), we also found no significant difference in average PSD-95 fluorescence across sessions (Figure 3E). Similar results were obtained when the gephyrin or PSD-95 fluorescence was normalized to the cell fill fluorescence (not shown).

### Impaired inhibitory synapse loss in amyloid pathology is not due to altered neuronal activity levels

Since structural inhibitory synapse loss in the dendritic shaft is an adaptation triggered by decreased experience-evoked neural activity (Chen et al., 2012; van Versendaal et al., 2012), its disruption in hAPP mice could be due to increased baseline spontaneous neuronal activity in these mice compared to WT. Using calcium imaging, we previously found that average spontaneous population activity (mean population dF/F0 of GCaMP6 transients) in the visual cortex recorded without visual experience is not significantly different in ∼5-6-month-old hAPP mice (Niraula et al., 2023). However, the same average population activity may arise despite differences in the distribution of activity patterns and correlational structure. We analyzed this data set of spontaneous activity over ∼228 seconds from 1020 WT and 885 hAPP neurons (Figure 4A). To assess whether the distribution of neurons exhibiting distinct mean calcium responses (quantified as dF/F0 (%)) during spontaneous activity differed, we categorized neurons into various bins based on their mean dF/F0 values. We found that the activity distribution is similar between the genotypes (Figure 4A).

**Figure 4.**
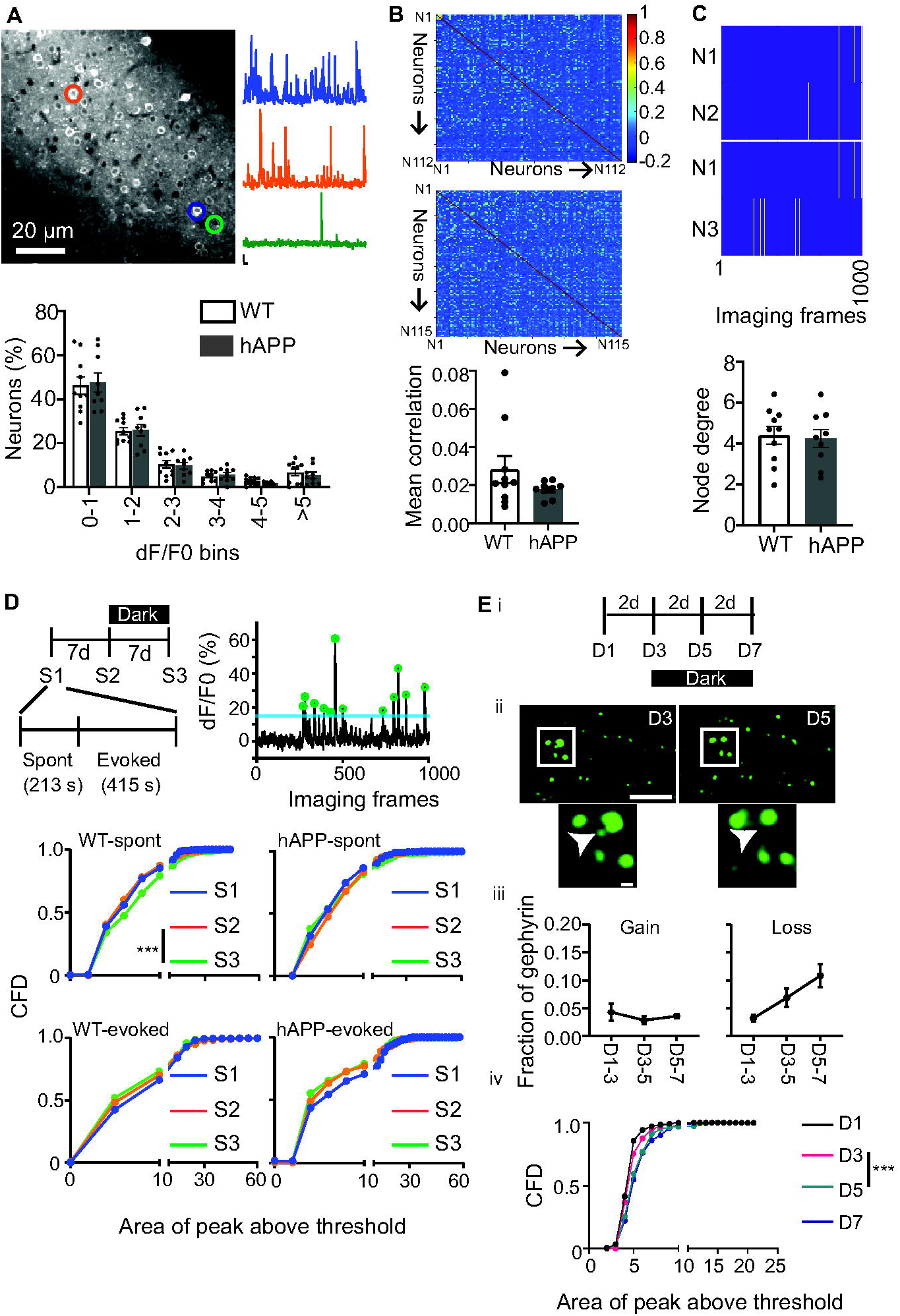
Baseline spontaneous activity is similar in WT and hAPP mice, whereas visual deprivation broadens it selectively in WT mice. **A.** Standard deviation projection image of a representative imaging field of neurons expressing GCaMP6s (top). Calcium transients (dF/F0) of three color-matched circled neurons with varying levels of activity. Percentage of neurons in different dF/F0 bins in WT and hAPP mice (bottom). **B**. Representative correlation matrices of WT and hAPP mice. (Bottom) Mean correlation from all mice. **C**. Representative images of two neurons that are functionally connected (top) or not connected (bottom) and the average node degree (right). n = 10 WT (1020 neurons) and 9 hAPP mice (885 neurons). Data are presented as mean ± SEM. Circles represent the average of each mouse. **D**. (Top left) Experimental paradigm - S1, S2, and S3 are imaging sessions separated by 7 days (7d). Between S2 and S3, mice were housed in darkness for one week. In each imaging session, both spontaneous (Spont) and visually evoked activities were imaged (indicated for S1). (Top right) Calcium transients during spontaneous imaging frames of an example neuron. The blue line represents the mean + 2 standard deviation threshold above which peaks (marked by green circles) were detected. (Middle-bottom) Cumulative frequency distribution (CFD) of average AUC per peak above the threshold for each neuron during spontaneous (middle) and evoked (bottom) activities in WT (left) and hAPP (right) mice. n = 390 (S1), 401 (S2), 419 (S3) WT neurons (5 WT mice) and 233 (S1), 206 (S2), and 209 (S3) hAPP neurons (4 hAPP mice). *** P<0.001 KS test. Alpha value was set to 0.025 to account for multiple (two – S1 vs. S2 and S2 vs. S3) comparisons (Bonferroni correction). The x-axis is modified for visualization purposes. **E.** (i) Timeline of gephyrin and jRGECO imaging sessions separated by two days (2d). Between day (D)3 and D7, mice were housed in complete darkness. (ii) Representative images of gephyrin puncta detected by GPHN-FingR-GFP on D3 and D5 (scale bar: 10 µm). The white box is magnified below, and the arrow points to a lost puncta (scale bar: 1 µm). (iii) Fraction of gephyrin puncta (total: 422 puncta) gained (left) and lost (right) between sessions. Gain did not significantly change over time, whereas loss increased (p<0.05, *F*_(1.69,8.46)_ = 5.8); repeated-measures ANOVA for loss. n = 6 WT mice). (iv) Cumulative frequency distribution of the average AUC per peak above the threshold of spontaneous calcium transients for each neuron. ***p<0.001 KS test. n = 140 (D1), 152 (D3), 165 (D5), and 122 (D7) neurons. Alpha value was set to 0.016 to account for multiple comparisons (D1 vs. D3, D3 vs. D5, D5 vs. D7; Bonferroni correction).

Changes to the correlational structure of neural activity may underlie impaired inhibitory synapse loss in hAPP mice. We obtained the mean correlation for each animal by averaging all the individual neuron-neuron correlations. We found that WT and hAPP mice had a similar average correlation (Figure 4B). The amplitude of individual calcium transients could be influenced by the number of coactive neurons (node degrees), even when the average calcium transient levels are similar between genotypes. We calculated the node degree of a neuron as the number of significant coactive neurons. Two neurons are considered significantly coactive if the number of imaging frames these neurons are coactive is greater than 95% of the cumulative distribution of coactivity generated by 1000 random circular shifts of activities between two neurons. Again, we found no difference in the average node degree of WT and hAPP mice (Figure 4C). Consistently, the mean AUC of individual calcium transients was also similar between the genotypes (WT: 7.5 ± 0.27, hAPP: 7.03 ± 0.19). These results indicate that impaired inhibitory synapse adaptation in hAPP mice may not be due to increased baseline spontaneous neuronal activity, raising the possibility that inhibitory synapses are not responsive to changes in experience-evoked activity in amyloidosis.

To test whether differential effects of visual deprivation on inhibitory synapse loss in WT and hAPP mice also manifest at the neural activity level, we imaged spontaneous and visually evoked calcium transients in the same mice over three imaging sessions, with each imaging session separated by one week. Mice were housed in complete darkness between the second and third sessions (Figure 4D). In each session, we imaged spontaneous and visually evoked calcium transients. Since inhibition regulates the temporal window of integration of synaptic excitation (Isaacson and Scanziani, 2011), we quantified the AUC of each calcium transient exceeding a threshold during the entire spontaneous imaging period (∼213 seconds). Similarly, we measured the AUC of calcium transients associated with visual stimuli (except gray screen) during the evoked imaging period (Figure 4D; Methods). We found that the average AUC per transient of each neuron did not significantly differ between the first two sessions for spontaneous and evoked activities for both genotypes, though there was a non-significant left shift in evoked activity in hAPP mice (Figure 4D middle and bottom). However, visual deprivation (between the second and third sessions) led to a significant right shift of average AUC per transient of spontaneous activity in WT but not hAPP mice (Figure 4D middle). Furthermore, visual deprivation did not significantly influence evoked AUC per transient in both genotypes (Figure 4D bottom). These results show that spontaneous neural activity, similar to inhibitory synapse loss, is elevated following one week of dark housing in WT but not hAPP mice. These results are also consistent with our recent findings that c-Fos expression in the visual cortex is increased in WT but not hAPP mice following one week of dark exposure (L’Esperance et al., 2023).

To further refine the time course of inhibitory synapse loss and increased spontaneous neuronal activity in WT mice, we labeled inhibitory synapses with a GPHN.FingR-GFP (Gross et al., 2013), an intrabody that recognizes gephyrin, and neurons with jRGECO, a red fluorescent calcium sensor (Dana et al., 2016). We imaged spontaneous activity and gephyrin puncta (not in the same neurons) every two days for 6 days, with mice housed in complete darkness for the last four days (Figure 4E). Between the first two sessions, when mice were housed in the normal light-dark cycle, the fraction of inhibitory synapses gained (0.04 ± 0.02) and lost (0.03 ± 0.01) were similar (Figure 4E). However, the loss was higher than the gain after two (gain: 0.03 ± 0.01, loss: 0.07 ± 0.02; p = 0.09, paired t-test) days in the dark but reached significance when tested after four days in the dark (gain: 0.04 ± 0.01, loss: 0.10 ± 0.02; p = 0.01, paired t-test; Figure 4E). We found that the broadening of spontaneous calcium transients occurred during the same period (two days of complete darkness), and no further broadening occurred during the next two days in darkness (Figure 4E).

### Disrupted clustering of excitatory and inhibitory synapse loss in amyloidosis

Though visual deprivation allowed us to identify the deficit in inhibitory synapse loss, whether amyloid pathology disrupts or alters compensatory inhibitory changes under non-perturbed conditions is unclear. The structural plasticity of excitatory and inhibitory synapses are spatially clustered (Chen et al., 2012). Therefore, if inhibitory synapses are not responsive to changes in excitatory synaptic activity, we reasoned that loss of excitatory synapses might not be associated with locally clustered inhibitory synapse loss in hAPP mice even under normal visual experience (12-hour light cycle). To assess this, we calculated the nearest neighbor distances of lost inhibitory synapses for every lost excitatory synapse under normal visual experience (Figure 5A). We found the median and mean nearest neighbor distances of lost inhibitory (target synapses) from a lost excitatory synapse (source synapses, where the source is the reference location to which target distances are measured) in WT mice to be 4 and 6.8 ± 1.2 (s.e.m) µm, respectively (Figure 5A). Interestingly, the distribution of nearest neighbor distances for hAPP mice was significantly right-shifted compared to WT mice, with the median and mean nearest neighbor distances of 7.7 and 13.2 ± 2.1 µm, respectively (Figure 5A). To assess whether the observed clustering distances are a reflection of altered density and number of lost synapses within a dendritic segment between genotypes, we randomized the distribution of the locations of synapses and their loss for each dendritic segment (Methods). Next, for the experimentally observed excitatory synapse loss (source) locations on the dendrite, we averaged 10,000 random nearest neighbor distances of inhibitory synapse loss (target) for each dendritic segment. We found that the observed mean for WT but not hAPP mice was significantly lower than random distribution (WT-randomized: 9.0 ± 1.3 µm; hAPP-randomized: 15.1 ± 2.2 µm). However, the median nearest neighbor distances of randomized distributions (WT-randomized: 8.9 ± 1.3 µm; hAPP-randomized: 14.9) were more right-shifted compared to observed medians for both genotypes, indicating local clustering is disrupted only for some excitatory synapse loss (source) in hAPP mice (Figure 5A).

**Figure 5.**
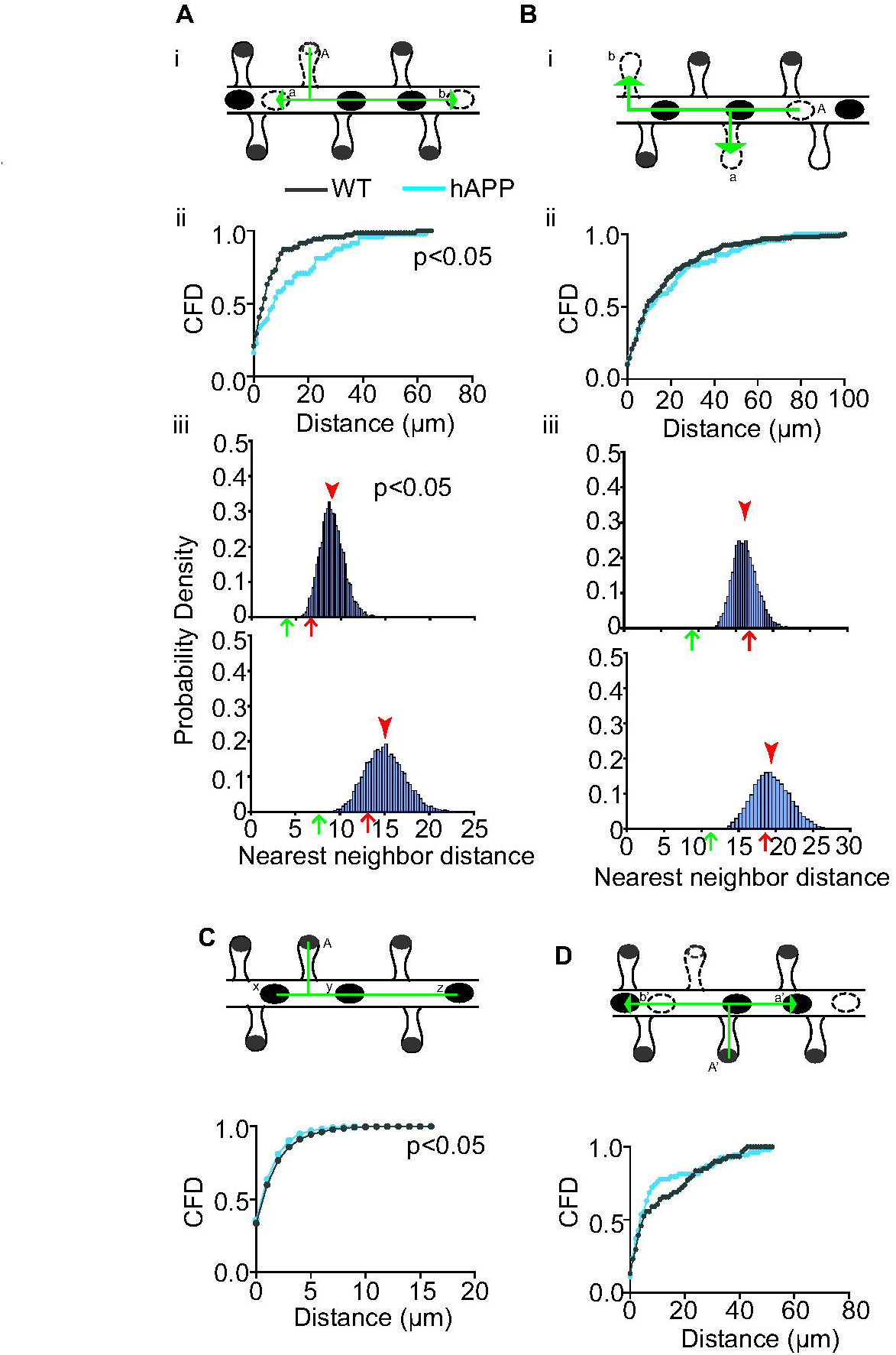
Local clustering of excitatory and inhibitory synapse loss is disrupted in hAPP mice. **A.** (i) A representative illustration of nearest neighbor analysis. “A” is an excitatory synapse that disappeared between imaging sessions. “a” and “b” are two inhibitory synapses that disappeared during the same interval. “a” will be the nearest inhibitory synapse loss (target) for the lost excitatory synapse “A” (source) in this dendritic segment. The distance to the nearest inhibitory synapse loss was calculated for every excitatory synapse lost. (ii) Cumulative frequency distribution (CFD) of nearest neighbor distances of inhibitory synapse loss to lost excitatory synapses. n = 71 (WT) and 48 (hAPP) lost excitatory synapses, P<0.05, Kolmogorov-Smirnov (KS) test. (iii) Distribution of average nearest neighbor distances of randomized inhibitory synapse loss to all experimentally observed locations of excitatory synapse loss from all dendrites in WT (top) and hAPP (bottom). Red arrowhead, red, and green arrows indicate the mean of random distribution, mean, and median observed values, respectively. p<0.05 (Z-test, left-sided). **B.** (i) “a” is the nearest excitatory synapse lost (target) to “A”, a lost inhibitory synapse (source). (ii) CFD of nearest neighbor distances of excitatory synapse loss to lost inhibitory synapses. n = 158 (WT) and 71 (hAPP) lost inhibitory synapses.(iii) Distribution of average nearest neighbor distances of randomized excitatory synapse loss to all experimentally observed locations of inhibitory synapse loss from all dendrites in WT (top) and hAPP (bottom). The red arrowhead indicates the mean of random distribution. **C.** (Top) “x” is the nearest inhibitory synapse (target) to “A,” the excitatory synapse (source). (Bottom) CFD of nearest neighbor distances of inhibitory synapses to all excitatory synapses. p<0.05 KS test, n = 2453 (WT) and 1851 (hAPP) excitatory synapses. **D.** (Top) a’ is the nearest gained inhibitory synapse (target) to A’, a gained excitatory synapse (source) between sessions. For every gained excitatory synapse, the distance to the nearest inhibitory synapse gain was calculated. (Bottom) CFD of nearest neighbor distances of gained inhibitory synapses (target) to all gained excitatory synapses (source). n = 61 (WT) and 54 (hAPP) gained excitatory synapses,

We next calculated whether similar changes to local clustering occur if inhibitory synapses are used as a source to identify nearest neighbor distances of target excitatory synapses. We found that the distribution of nearest neighbor distances of WT and hAPP mice did not differ, and their mean was not significantly different from the random distribution (Figure 5B). However, the median of both genotypes was lower than the random distribution (median: 9.4 µm (WT), 16.1 µm (WT-randomized), 11 µm (hAPP), 19.2 (hAPP-randomized), indicating not all excitatory synapse loss (target) in relation to lost inhibitory synapse (source) are random.

To further confirm that the closer clustering of inhibitory synapse loss to lost excitatory synapses in WT compared to hAPP mice is not a reflection of spacing between excitatory and inhibitory synapses in these genotypes, we measured the nearest neighbor distances of inhibitory synapses (target) to every excitatory synapse (source). We found they were very similar (median WT: 1.04 µm and hAPP: 0.96 µm), though there was a small but significant left shift for hAPP mice (Figure 5C). Furthermore, the distribution of nearest neighbor distances of lost excitatory synapses (mean: 14.8 ± 2.2 (WT), 17.9 ± 3.2 (hAPP); median: 8.9 (WT), 10.3 (hAPP)) or lost inhibitory synapses (mean: 8.6 ± 0.8 (WT), 10.1 ± 0.9 (hAPP); median: 4.4 (WT), 6.0 (hAPP)) with each other (target and source synapse type are same) were not significantly different. We also did not observe a significant difference between the genotypes when we compared the distribution of the nearest neighbor distances of gain of inhibitory synapses (target) in relation to source excitatory synapse gain (Figure 5D), and both genotypes showed close clustering (median: 5 µm (WT), 4 µm (hAPP)). Together, these results show that the right shift in the distribution of the nearest neighbor distances of lost excitatory and inhibitory synapses in hAPP mice represents an impairment in their local clustering.

## Discussion

Direct visualization of synaptic proteins *in vivo*, repeated imaging of the same synapses, and the use of the visual cortex as a model system, which is amenable to altering experience-evoked activity, allowed us to unearth a pathological feature of amyloid that would not be directly evident by visualizing the density or baseline dynamics of synapses either *in vitro* or *in vivo*. Multiple animal studies have uncovered the vulnerability of the visual cortex to AD-related pathology and subtle synaptic dysfunction without overt degeneration could contribute to visual deficits observed in a subset of AD patients (Armstrong, 1996; Grienberger et al., 2012; William et al., 2012; Liebscher et al., 2016; Korzhova et al., 2021; William et al., 2021; Kurucu et al., 2022; Papanikolaou et al., 2022; Niraula et al., 2023).

In amyloid pathology, we identified a selective disruption in the experience-dependent structural adaptation of inhibitory synapses on the dendritic shaft. The density or baseline structural dynamics of inhibitory synapses were not significantly perturbed. Furthermore, the baseline structural dynamics of excitatory synapses and their response to altered experience remain unchanged in the same neurons. These results indicate that structural plasticity deficits in the inhibitory system emerge before impairments in the structural plasticity of excitatory synapses in amyloidosis.

Hyper- and hypoactivity of neurons and brain regions in Alzheimer’s disease are thought to arise due to defective homeostatic adaptation (Jang and Chung, 2016; Frere and Slutsky, 2018; Styr and Slutsky, 2018). Excitatory synapse loss is thought to be one of the maladaptive plasticity mechanisms to restrain hyperactivity and could lead to hypoactivity when unchecked (Styr and Slutsky, 2018). A reduction in excitatory synapse density and brain activity is observed in the later stages of AD (Terry et al., 1991; DeKosky, Scheff and Styren, 1996; Dickerson et al., 2005; Celone et al., 2006; Scheff and Price, 2006), indicating disrupted inhibitory plasticity mechanisms may underlie lack of compensation to reduced neural activity associated with excitatory synapse loss. Though the order of emergence of hyper- and hypoactivity is still unclear, they must closely follow each other to maintain average activity levels as the total energy budget of the brain is fixed. Consistently, hyper- and hypoactive neurons are present together in mouse models of amyloidosis (Busche et al., 2008; Grienberger et al., 2012; Rudinskiy et al., 2012; Niraula et al., 2023). We recently observed that the increase in neuronal hyper- and hypoactivity occurs at a stage where postsynaptic densities of excitatory and inhibitory synapses are comparable to nonpathological controls (Niraula et al., 2023). Therefore, mechanisms other than excitatory synapse loss, such as changes to the size, stability, and physiology of inhibitory synapses, may contribute to hypoactivity. Consistently, gephyrin levels were found to be higher in amyloid pathology (Hales et al., 2013; Kiss et al., 2016), though the literature is inconsistent on the direction of change in the GABAergic system concerning favoring or opposing hyperactivity. The insensitivity of inhibitory synapses to activity perturbation and local decoupling of excitatory and inhibitory synapse loss indicates that inhibitory synapses in amyloid pathology may not compensate for the reduction in activity levels.

In line with the role of inhibitory synapse plasticity in maintaining neuronal activity homeostasis, synaptic inhibition is increased in the light phase of the light-dark cycle to compensate for increased excitability in the dark phase (Bridi et al., 2020). Consistently, neurons in the visual cortex do not exhibit significant activity changes across light-dark cycles (Torrado Pacheco et al., 2019). However, two-day dark exposure elicits disinhibition (Huang et al., 2015), and increased spontaneous activity has been observed after one week (Bridi et al., 2018). We also observed that the AUC of calcium transients increased following dark exposure in WT but not hAPP mice, which also do not exhibit increased inhibitory synapse loss. Intriguingly, we did not observe an increase in the AUC of calcium transients for visually evoked activity, indicating inhibitory synapse loss associated with long-term dark exposure may facilitate cross-modal inputs. In this view, inhibitory synapse loss may not serve a homeostatic role and may promote cross-modal plasticity by increasing the temporal window of synaptic activity during each calcium transient. Since dendritic disinhibition promotes learning-associated plasticity (Letzkus, Wolff and Luthi, 2015; Mohler and Rudolph, 2017; Artinian and Lacaille, 2018; Wu, Miehl and Gjorgjieva, 2022), disrupted inhibitory synapse adaptation may also interfere with learning or cross-modal plasticity in amyloidosis. Though the direction of neural activity and inhibitory synaptic plasticity in both genotypes are consistent, it is challenging to extrapolate the small effect size of structural synaptic changes to neural and microcircuit activity.

Disrupted inhibitory synapse loss following visual deprivation in hAPP mice may arise due to multiple mechanisms. There may be no intrinsic defect in inhibitory synapses that prevents them from undergoing synapse loss. For instance, an elevated baseline activity may not necessitate inhibitory synapse loss in hAPP mice. However, our spontaneous activity analyses indicate that baseline spontaneous activity (without visual experience) in the visual cortex of hAPP mice at this age is similar to nonpathological controls based on multiple metrics. Therefore, impaired inhibitory synapse adaptation is unlikely due to higher spontaneous activity in hAPP mice in the dark. Alternatively, the neurons in hAPP mice may lack sensors to detect changes in activity. We found that excitatory synapses respond to visual deprivation by reducing ongoing spine gain in the neurons exhibiting inhibitory synapse loss deficits. This argues against a general defect in sensing reduction in activity levels. We propose that at least a fraction of inhibitory synapses in neurons of hAPP mice become disengaged from changes in excitatory activity. As a result, these synapses are more stabilized, and local alteration in an excitatory activity does not destabilize them.

Is inhibitory synapse loss a response to or a cause of excitatory synapse loss? The clustered loss of excitatory and inhibitory synapses is disrupted in hAPP mice only when the proximity of inhibitory synapse loss (target) to lost excitatory synapses (source) was measured. The distance of the nearest excitatory synapse loss (target) to lost inhibitory synapses (source) is larger and did not differ between hAPP and WT mice. These results suggest that inhibitory synapses in amyloidosis are deficient in sensing or responding to changes in excitatory activity. The mechanisms that drive inhibitory synapse loss following excitatory synapse loss are unclear. Multiple signaling pathways have been shown to enhance or weaken inhibitory synapses following enhancement in excitation (Rutherford et al., 1997; Swanwick, Murthy and Kapur, 2006; Chen et al., 2012; Flores et al., 2015; Hu et al., 2019; Ravasenga et al., 2022). It is unclear whether these signaling mechanisms also weaken inhibitory synapses following the weakening of excitatory synapses, possibly by maintaining their co-stability. The direction of clustering of excitatory synapse and inhibitory synapse gain or loss may differ depending on the context. In the learning context, disinhibition is favorable, and consistently, an increase in excitatory synaptic strength was shown to weaken inhibitory synaptic strength within 3 µm (Ravasenga et al., 2022). Our study differs from this study since we looked at clustering of excitatory and inhibitory synapse loss, but interestingly, the median distance (4 µm) of clustering was very similar in WT mice. The reduction in local clustering of excitatory and inhibitory synapse loss may be a consequence of the increased stability of gephyrin from degradation. Multiple posttranslational modifications, particularly phosphorylation, influence gephyrin stability (Tyagarajan et al., 2011; Kuhse et al., 2012; Kalbouneh et al., 2014; Zacchi, Antonelli and Cherubini, 2014; Flores et al., 2015; Wang et al., 2015; Zhou et al., 2021). Kinases that phosphorylate gephyrin and alter its stability, such as CDK5, are dysregulated in AD (Kiss et al., 2020). Whether CDK5-mediated phosphorylation or other posttranslational mechanisms are involved in disrupted disinhibition remains to be tested.

Visual deprivation has been shown to elicit rapid disinhibition due to reduced activity of parvalbumin-expressing interneurons (Hengen et al., 2013; Kuhlman et al., 2013; Barnes et al., 2015). Parvalbumin-expressing neurons primarily target perisomatic regions and proximal dendrites, whereas somatostatin-expressing interneurons target distal dendrites (Di Cristo et al., 2004). However, their localization is not mutually exclusive. We did not find a spatially restricted dendritic disinhibition, and the reduction of synapse density by deprivation appeared to be more uniform across the length of the dendrite. Future work with cell-type specific labeling is required to test whether structural disinhibition happens at both parvalbumin and somatostatin innervations.

## Author contributions

SN performed the experiments. SN and JS performed data analyses. SSY and JS initiated the concept and designed the experiments. JS wrote the paper and supervised the research.

